# Striatal dopamine availability peaks in adulthood and undergoes age-dependent modulation

**DOI:** 10.64898/2026.11.10.765699

**Authors:** Samantha R. White, J. Hoon Shin, Hannah C. Goldbach, Marta Andujar, Evan S. Swanson, Rachele Rimondini, Abigail L. Carr, Lucy G. Anderson, Sydney E. Cerveny, Peyton Q. Harmon, Anne Claire Tangen, Kenneth A. Pelkey, Reza Azadi, Alexander C. Cummins, Christopher J. McBain, Bruno B. Averbeck, Veronica A. Alvarez

## Abstract

Age is a risk factor for several psychiatric disorders. An adolescent peak in striatal dopamine availability is thought to contribute to the neuropsychiatric vulnerability. However, the lifespan trajectory of striatal dopamine release capacity is undefined in most species. Here, we measured lifespan trajectories of striatal dopamine in primates and rodents and identified a delayed developmental peak in adulthood and age-dependent changes in dopamine release regulation by GABA and acetylcholine across species.

## MAIN TEXT

Neuronal identities are largely specified in early development (Silbereis et al., 2016). However, there are substantial ongoing brain changes across the lifespan (Alldritt et al., 2024; Bethlehem et al., 2022; Hammelrath et al., 2016). One notable example is the considerable structural and functional reorganization of fronto-striatal circuitry during adolescence (Bo et al., 2014; Choi et al., 2023). Within the fronto-striatal circuitry, dopamine supports the remodeling through its regulation of synaptic plasticity, excitability, and neurotransmitter release (Lieberman et al., 2017; Kozorovitskiy et al., 2015). The circuit remodeling also coincides with increasing dopaminergic innervation throughout adolescence, as shown by anatomical studies in primates and rodents (Rosenberg and Lewis, 1995; Avramescu et al., 2024). Further, changes in dopamine receptors and transporter expression were demonstrated over the lifespan in multiple species (Karrer et al., 2017; Johansson et al., 2023; Lai et al., 1987; Meng et al., 1999). This evidence has led to the proposal that the heightened reward sensitivity seen during adolescence may be driven by a transient peak in dopamine availability that precedes the maturation of prefrontal inhibitory control (Casey et al., 2008; Wahlstrom et al., 2010). However, indirect measures of striatal dopamine availability suggest a steady rise during adolescence with a peak in young adulthood (Larsen et al., 2020; Parr et al., 2022), later than predicted by behavioral and anatomical models. Whether the lifespan trajectory of dopamine release capacity aligns with these indirect measures or traditional models has not been shown.

In addition to dopamine availability, the mechanisms that locally govern striatal dopamine release may also change across the lifespan. Recent data from primates and rodents have demonstrated striatal dopamine release is highly regulated by GABA and acetylcholine (Lopes et al., 2019; Kramer et al., 2020; Liu et al., 2022; Shin et al., 2025; Zhang et al., 2025) and these mechanisms can operate independently of midbrain dopamine neuron activity to shape plasticity and striatal output (Mohebi et al., 2019; Eshel et al., 2024). Thus, the maturation of the GABA and acetylcholine regulations of dopamine release may serve as candidate mechanisms that improve cognitive control from early adolescence to adulthood.

Here we determine the lifespan trajectory of dopamine availability and local regulation by GABA and acetylcholine in the striatum of macaques and mice. Defining the normative trajectory is critical for improving mechanistic understanding of developmental brain changes and sensitive periods, ultimately relevant for age-dependent vulnerability to psychiatric disorders (Andersen, 2003; Solmi et al., 2022; Ulhaas et al., 2023).

To examine cross-species striatal dopamine signaling over the lifespan, we used fast scan cyclic voltammetry (FSCV) to measure electrically evoked dopamine release and clearance in acute brain slices from male and female macaques (*Macaca mulatta,* n = 22) and mice (*Mus musculus*; n = 59) spanning approximately 70% of the lifespan for each species (macaque: 2-17 years old; mice: 3-70 weeks; **Extended Data Fig 1a**; Osigo et al., 2025; Simmons, 2016). We focused our recordings in the dorsal striatum as it is a conserved brain region with major cortico-thalamic convergence and established roles in reward sensitivity, action selection, motor control, and reinforcement learning (Balleine et al., 2007; McFarland and Haber, 2000; Pan et al., 2010).

We first examined the amplitude of the evoked dopamine signals as a function of stimulation intensity (**Extended Data Fig 1b**). To assess maximal dopamine release capacity (‘evoked dopamine’) we analyzed the response elicited by the highest stimulation intensity (400 μA; **Extended Data Fig 1b).** Across age, evoked dopamine followed an inverted-U curved trajectory in both macaques (*effective degrees of freedom (edf)* = 2.42; Generalized Additive Model w/ penalized spline; **Methods**) and mice (*edf* = 2.55; **Fig 1a**). In macaques, evoked dopamine increased from 2 years to an estimated peak at 7.1 years before declining modestly through the oldest measured timepoint at 17 years (**Fig 1a**). In mice, evoked dopamine increased from 3 weeks to an estimated peak at 29.6 weeks before declining towards 58 weeks (**Fig 1a**). To account for differences in longevity and compare trajectories across species, we normalized age to average lifespan in captivity. Despite marked species differences in release magnitude (mean ± s.e.m.: macaque: 553 ± 23 nM v. mouse: 835 ± 27 nM), lifespan-normalized trajectories peaked at nearly identical relative ages (macaque, 28.52%; mouse, 28.51%) and did not differ in shape (*edf* = 1.01; **Fig 1b**), supporting a broadly conserved chronological trajectory of dopamine release in representative primate and rodent species.

**Fig. 1.**
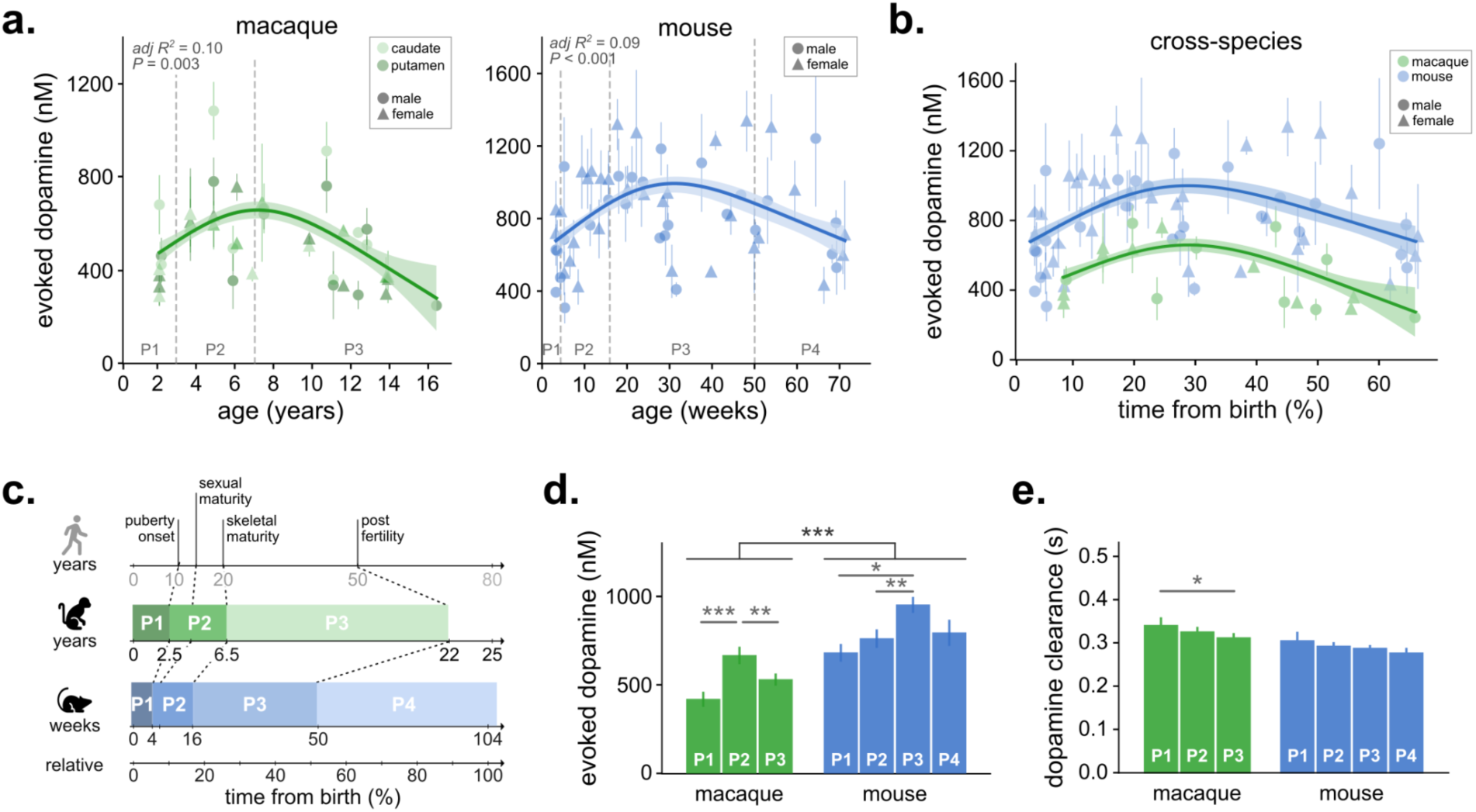
Evoked dopamine and clearance in the striatum follow similar lifespan trajectories in macaques and mice with species-specific developmental timing. a, Maximum electrically evoked striatal dopamine release plotted against age in macaques (left; N = 22 animals, n = 124 slices) and mice (right; N = 58, n = 189). Species-specific generalized additive models (GAMs; thin-plate regression splines, k = 4) revealed nonlinear relationships between age and evoked dopamine in macaques (F = 5.50, edf = 2.43, P = 0.003; 11.7% deviance explained) and mice (F = 7.48, edf = 2.56, P < 0.001; 10.3% deviance explained). In macaques, light and dark green denote caudate and putamen, respectively; circles and triangles denote males and females in both species. Shading denotes ±s.e.m. b, Evoked dopamine plotted against age normalized to average species lifespan. A GAM with penalized splines revealed a nonlinear association between relative age and dopamine release (edf = 5.47, χ² = 57.46, P < 0.001). Dopamine release was greater overall in mice (blue) than macaques (green; β = 350.87, z = 9.93, P < 0.001), whereas the species-specific age smooth was not significant (edf = 1.01, χ² = 0.30, P = 0.595), providing no evidence that the shape of the lifespan trajectory differed between species after accounting for overall release magnitude. Residual variance was greater in mice (β = 0.395, z = 4.79, P < 0.001). c, Cross-species developmental and aging milestones used to define lifespan periods. Sampled ages spanned approximately 70% of the average lifespan in macaques (2 - 17 years) and mice (3 - 68 weeks). d, Evoked dopamine release across lifespan periods. Release differed by period in macaques (F_2,117_ = 8.97, P < 0.001) and mice (F_3,181_ = 4.45, P = 0.005), with an overall effect of species (F_1,310_ = 53.62, P < 0.001) and a Period × Species interaction (F_2,279_ = 7.68, P < 0.001). In macaques, release was greater in P2 than in P1 (t_119_ = 4.23, P < 0.001) or P3 (t_119_ = 3.68, P = 0.001). In mice, release was greater in P3 than in P2 (t_184_ = −3.26, P = 0.007) or P1 (t_184_ = 2.60, P = 0.049). e, Dopamine transient decay time constant (τ) across lifespan periods. τ did not differ overall between species (F_1,312_ = 1.77, P = 0.18). In macaques, τ showed a trend across lifespan periods (F_2,122_ = 2.99, P = 0.054) and was lower in P3 than P1 (t_124_ = −2.40, P = 0.046), consistent with faster dopamine clearance at the later period. τ did not differ across lifespan periods in mice (F_3,180_ = 0.84, P = 0.47). *Repeated-measures ANOVA was used unless otherwise indicated. Post hoc comparisons were performed using estimated marginal means with Tukey adjustment for multiple comparisons. Error bars represent mean ± s.e.m. N, animals; n, slices. *P < 0.05, **P < 0.01, ***P < 0.001.*

We next analyzed the same data relative to conserved biological milestones. Because developmental windows differ in timing and duration across species, we defined lifespan periods using puberty onset, skeletal maturation, and onset of reproductive decline as boundaries. This yielded three lifespan periods in macaques and mice: P1-birth to puberty onset (macaque: 2.5y; mouse: 4w; Resko et al., 1982; Chester et al 2026); P2 - puberty onset to skeletal maturation (macaque: 6.5y; mouse: 16w; Trotter et al., 1975; Jilka 2013); and P3 - skeletal maturation to onset of reproductive decline (macaque: 20y; mouse: 50w; Walker and Herndon, 2008; Finch 2014). A fourth post-reproductive period (P4) was only available in mice (**Fig 1c**).

In macaques, evoked dopamine was largest in the post-pubertal P2 period (664 ± 44 nM) compared to the earlier (P1: 417 ± 38 nM) and later lifespan periods (P3: 528 ± 29 nM; **Fig 1d**). By contrast, mice showed largest evoked dopamine in the post-maturity period P3 (948 ± 40nM) compared to P1 (678 ± 45 nM), P2 (758 ± 47 nM) and P4 (791 ± 69nM; **Fig 1d**). A significant species x period interaction indicated that evoked dopamine release peaked earlier relative to developmental milestones in macaques despite parallel lifespan-normalized trajectories (**Fig 1d**). Release was also greater in male than female macaques and modestly greater in caudate than putamen, although neither effect interacted with lifespan period **(Extended Data Fig. 1e**). Mice showed no sex difference across periods (**Extended Data Fig 1f**). Together these results support lower overall striatal evoked dopamine in primates than rodents with parallel curvature (Shin et al., 2025; Calipari et al., 2012), while revealing species-specific developmental timing of striatal dopamine release capacity.

Because striatal dopamine signals reflect both midbrain dopamine firing and local terminal regulation (Liu and Kaeser, 2019; Holly et al., 2025) we next examined mechanisms that could contribute to these lifespan trajectories. Dopamine clearance, largely driven by dopamine transporters (DAT) in the striatum (Sulzer et al., 2016), was assessed from the decay time constant of evoked transients. Based on reported age-related declines in DAT expression (Salvatore et al., 2003; Harada et al., 2002), we predicted that clearance would slow with age. Mean decay time constants were similar in macaques (0.31 ± 0.01 s) and mice (0.30 ± 0.01 s; **Fig 1e**), but developmental trends differed. Contrary to our prediction, clearance became faster with age in macaques with decay time constants decreasing from P1 (0.34 ± 0.02 s) to P3 (P3: 0.29 ± 0.01 s; **Fig 1e**) whereas mice remained stable across periods (**Fig 1e**). No sex effects were detected (**Extended Data Table**). These findings suggest that age-related changes in clearance cannot be explained by DAT expression alone and may reflect changes in transporter function or other terminal properties.

It has also been established that intra-striatal GABA and acetylcholine signaling regulate dopamine release and differ substantially across species (Lopes et al., 2019; Kramer et al., 2020; Liu et al., 2022; Shin et al., 2025; Zhang et al., 2025). GABA and nicotinic acetylcholine receptor (GABAR and nAChR) densities also change dynamically across development in rodents (see Thorpe et al., 2019). Adult macaques show greater GABA-mediated inhibition of dopamine release, whereas mice show greater acetylcholine-mediated facilitation (Shin et al., 2025), prompting us to ask whether these mechanisms also follow distinct developmental trajectories. Bath application of GABA_A_ and GABA_B_ receptor antagonists potentiated evoked dopamine release in macaques (+51 ± 4%) and, to a lesser extent, in mice (+12 ± 2%; **Fig. 2a,b**). Antagonist-induced potentiation increased with age (**Extended Data Fig. 2a**) and lifespan period (**Fig. 2b**) in both species. In macaques, it increased from P1 (+32 ± 8%) through P2 (+50 ± 6%) to P3 (+62 ± 5%). In mice, potentiation was absent in P1 (−4 ± 4%) but emerged after puberty and increased through P2 (+13 ± 3%), P3 (+14 ± 3%) and P4 (+18 ± 5%). These effects did not differ by sex or, in macaques, region (**Extended Data Table**). Thus, GABAergic regulation is substantially greater in macaques but strengthens with maturation in both species.

**Fig. 2.**
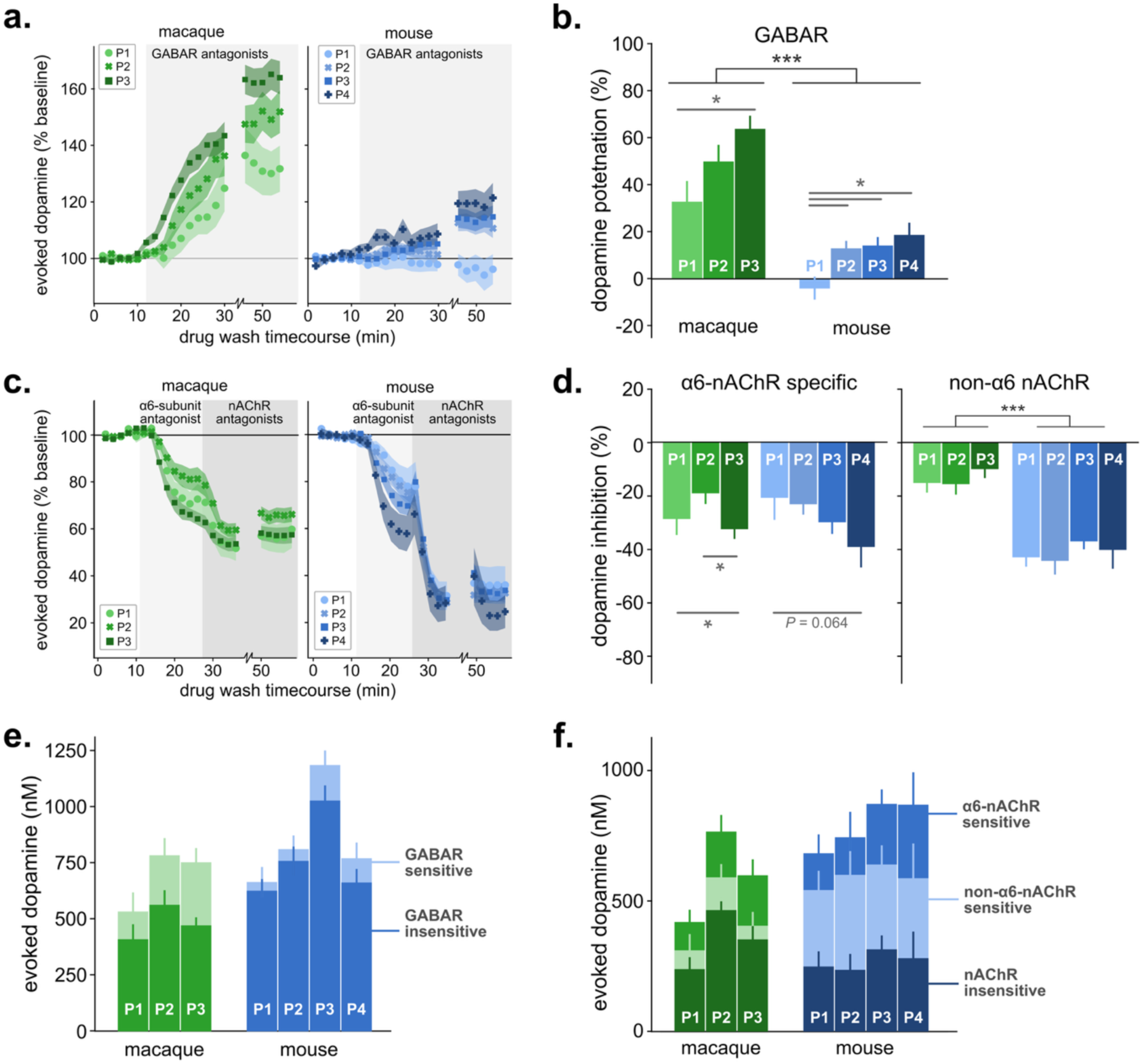
GABAergic and cholinergic regulation of evoked dopamine signals change throughout the lifespan in macaque and mouse. a, Time course of evoked dopamine potentiation following GABA receptor antagonism in macaque and mouse striatal slices. Peak dopamine amplitude was normalized to pre-drug baseline. Symbols and shading show mean ± s.e.m. for slices from each lifespan period (macaque: *N* = 19, n = 57; mouse: N = 40, n = 78). Gray shading indicates drug application. b, Mean potentiation of evoked dopamine by GABA receptor antagonists across lifespan periods. Potentiation differed by species (F_1,130_ = 98.93, P < 0.001) and lifespan period (F_1,130_ = 5.87, P = 0.0008), with no period × species interaction (F_2,118_ = 1.11, P = 0.33). Species-specific analyses showed effects of lifespan period in macaques (F_2,50_ = 4.62, P = 0.014) and mice (F_3,70_ = 3.28, P = 0.026). In macaques, potentiation was lower in P1 than P3 (t_52_ = 2.64, P = 0.029). In mice, P1 was lower than P2 (t_73_ = 2.63, P = 0.050), P3 (t_73_ = 2.62, P = 0.052) and P4 (t_73_ = −2.82, P = 0.031). c, Time course of evoked dopamine inhibition following sequential application of the α6-nAChR antagonist α-conotoxin PIA (0.1 μM; light gray) and DHβE (1 μM; dark gray). Peak dopamine amplitude was normalized to pre-drug baseline. Symbols and shading show mean ± s.e.m. for slices from each lifespan period (macaque: N = 16, n = 56; mouse: N = 43, n = 89). d, Mean inhibition of evoked dopamine by α6-containing nAChRs (left) and the remaining non-α6 nAChR component (right) across lifespan periods. α6-nAChR- mediated inhibition did not differ by species (F_1,137_ = 0.18, P = 0.67) but varied by lifespan period (F_3,137_ = 3.94, P = 0.0097). In macaques, inhibition differed across periods (F_2,26_ = 5.33, P = 0.008), with greater inhibition in P3 than P2 (t_46_ = 3.26, P = 0.006). In mice, the period effect did not reach significance (F_3,81_ = 2.52, P = 0.064). The non-α6 nAChR component differed by species (F_1,125_ = 75.72, P < 0.001) and lifespan period (F_3,125_ = 2.87, P = 0.040), although species-specific analyses did not detect a period effect in macaques (F_2,45_ = 1.35, P = 0.26) or mice (F_3,70_ = 0.78, P = 0.509). e, Evoked dopamine release before and after GABA receptor antagonism across lifespan periods. Saturated colors denote baseline evoked dopamine and lighter colors denote responses after 30 min of antagonist application. In macaques, the GABA receptor antagonist effect did not differ by region (F_1,50_ = 0.73, P = 0.39) or sex (F_1,50_ = 1.52, P = 0.22). No sex effect was detected in mice (F_1,70_ = 2.80, P = 0.09). f, Evoked dopamine release before and after sequential nAChR blockade across lifespan periods. Saturated colors denote baseline evoked dopamine, lighter tones denote α6-nAChR-insensitive release after α-conotoxin PIA, and darker tones denote nAChR-insensitive release after subsequent DHβE application. *Repeated-measures ANOVA was used unless otherwise indicated. Post hoc comparisons were performed using estimated marginal means with Tukey adjustment for multiple comparisons. Error bars and shaded areas represent mean ± s.e.m. N, animals; n, slices. *P<0.05, **P<0.01, ***P<0.001*

We next examined developmental changes in cholinergic regulation of striatal dopamine release. We focused first on α6-containing nAChRs due to their known relevance in cholinergic regulation of striatal dopamine in macaques (Zhang et al., 2025; Quik et al., 2011). We hypothesized that increasing nAChR-dependent facilitation of dopamine might contribute to the developmental increase in evoked dopamine (Zoli et al., 2002; Grady et al., 2007; Han et al., 2000; Reynolds et al., 2025; Shin et al., 2025). Bath application of α-conotoxin PIA (0.1 µM), which selectively blocks α6-containing nAChRs, reduced evoked dopamine release similarly in macaques (27 ± 2%) and mice (28 ± 2%; Fig. 2d). This suppression was greater in female than male macaques (M: 23 ± 3%; F: 30 ± 3%; **Extended Data Fig. 2c**); but did not differ by sex in mice (M: 25 ± 4%; F: 31 ± 3%; **Extended Data Fig. 2d**). Across lifespan periods, α6-dependent regulation differed between species. In macaques, antagonist-induced suppression was smallest during P2 (P1: 28 ± 5%; P2: 18 ± 4%; P3: 32 ± 3%; **Fig. 2d**), whereas in mice, it increased progressively from P1 (20 ± 7%) through P2 (23 ± 3%), P3 (29 ± 4%) and P4 (41 ± 7%; **Fig. 2d; Extended Data Fig. 2c**).

Subsequent blockade of non-α6 containing nAChRs with high concentration DHβE (1 µM; Kirsch et al., 2016) produced additional suppression of dopamine release in both species, with a larger effect in mice (40 ± 2%) than macaques (13 ± 2%; **Fig. 2d**). In macaques, this DHβE-sensitive component also differed by sex (M: 15 ± 2%; F: 10 ± 2%) and region (caudate: 16 ± 2%; putamen: 8 ± 2%; **Extended Data Fig. 2d**). Unlike the α6-sensitive component, the additional DHβE-sensitive component remained relatively stable across lifespan periods in macaques (P1: 15 ± 3%; P2: 15 ± 3%; P3: 10 ± 3%) and mice (P1: 42 ± 3%; P2: 44 ± 4%; P3: 37 ± 2%; P4: 40 ± 6%; **Fig. 2d**). Overall nAChR-dependent regulation was therefore smaller in macaques (40 ± 2%) than mice (67 ± 2%), consistent with previous findings (Shin et al., 2025). In mice, increasing α6-nAChR-dependent regulation paralleled the developmental increase in evoked dopamine, raising the possibility that α6-dependent cholinergic facilitation contributes to the later maturation of dopamine release. In macaques, by contrast, peak P2 release coincided with the smallest α6-dependent component, arguing against enhanced α6-mediated facilitation as the mechanism underlying this peak.

In summary, this systematic cross-species characterization revealed parallel, inverted U-shaped lifespan trajectories of striatal dopamine availability in macaques and mice. In both species, dopamine release capacity peaked at approximately 30% of the lifespan, corresponding to a post-adolescent young-adulthood period. Alignment to conserved biological milestones revealed this peak coincides with skeletal maturity in macaques and was further delayed in mice. In either case, this timing does not support a model in which an adolescent peak in dopamine drives the heightened reward sensitivity and impulsivity observed in adolescence (Wasserman et al., 2021; Wahlstrom et al., 2010). Instead, our findings support the idea that the ongoing maturation of striatal dopamine function continues beyond adolescence into adulthood, consistent with human imaging studies showing that dopamine-related measures peak in young adulthood (Larsen et al., 2020; Parr et al., 2022). This prolonged maturation suggests that the sensitive period may extend later than previously anticipated (Semple et al., 2013; Martin and Cook, 2014; Marin, 2016), providing a longer window during which experiences and environmental factors can have lasting effects on circuit architecture.

The regulation of dopamine release also changes across the lifespan. GABA mediated inhibition of dopamine release strengthened with age in both species. The underlying mechanism of this age-dependent increase in inhibition may include more GABA content and/or stronger receptor expression and efficacy. Further characterization of the striatal GABA neurons across the lifespan would provide key insights. The strong age-dependent restraint of dopamine by intra-striatal GABA may contribute to the dopamine-related pathophysiology underlying some psychiatric disorders. For example, disruptions in GABA regulation of dopamine across the lifespan may contribute to hypodopaminergic states in depression and hyperdopaminergic states in psychotic disorders (Srivastav et al., 2023; Syndor et al., 2025).

In contrast to GABA, the cholinergic facilitation of dopamine release changes with age differently in macaques and mice. We predicted that the cholinergic facilitation may be maximal at the highest dopamine availability, contributing to its maximum. However, in macaques, the cholinergic facilitation of dopamine was the smallest during the post-pubertal period, arguing against the cholinergic facilitation driving the dopamine peak. In mice, cholinergic facilitation increased with age and was overall larger than in macaques. The cholinergic modulation continues to increase past the adulthood peak of dopamine, thus arguing against the cholinergic facilitation may contribute to the peak emergence. Together these results suggest the developmental maxima in dopamine release emerges from multiple regulatory processes rather than a single local mechanism. Furthermore, the relative contribution of these mechanisms across age differed between species as described below.

We also found that dopamine transients decayed more rapidly in older compared to younger macaques but remained relatively stable in mice. Faster clearance improves the temporal precision of dopamine signals and may contribute to the refinement of both motor and cognitive control observed in young adulthood (Bu et al., 2021; Vaughan and Foster, 2013). These results suggest DAT activity shifts across the lifespan and is part of frontostriatal circuit maturation.

We identified several species differences independent of age. Dopamine release was overall larger in mice compared to macaques, in part due to species difference in dopamine release regulation. These findings are consistent with prior comparative work done in only adult subjects (Shin et al., 2025, Calipari et al., 2012). In macaques, we found an overall stronger GABA mediated suppression of dopamine release. In mice, we found an overall stronger cholinergic facilitation of dopamine. In fact, blocking GABA receptors minimized the magnitude difference between macaque and mouse and blocking nicotinic receptors (α6-nACh) resulted in similar dopamine amplitudes across the species. Thus, the magnitude and the regulation of striatal dopamine availability is different between macaques and mice.

A consideration of these *ex vivo* recordings is the local electrical stimulation that nonspecifically activates nearby axons and neurons, including the dopamine neuron axons within the striatum. The resulting dopamine signals therefore reflect dopamine terminal function and local regulation, and unlike *in vivo* dopamine dynamics, dopamine signals are not spontaneous or behaviorally driven in response to dopamine cell body firing. Evoked signals in these *ex vivo* preparations reflect both direct stimulation of dopamine axons and indirect intra-striatal mechanisms involving other local axons and neurons, including GABAergic and cholinergic cells. We use receptor antagonists to probe the contribution of GABA and acetylcholine to these direct and indirect mechanisms, targeting GABA and nicotinic acetylcholine receptors because they are expressed on dopamine axons (Lopes et al., 2019; Kramer et al., 2020; Liu et al., 2022; Shin et al., 2025; Zhang et al., 2025; Quik et al., 2011).

These reported developmental changes in dopamine availability and release regulation may provide new avenues for understanding neuropsychiatric vulnerability. Prefrontal cortical inputs regulate striatal dopamine release in part through cholinergic interneurons and acetylcholine facilitation of axonal dopamine release (Shin et al., 2020; Goldbach et al., 2026). Maturation of these cortical projections and of the cholinergic modulation may contribute to the age-dependent improvements in cortical control over behavior from childhood to adulthood. Open questions remain about age-dependent changes in other mechanisms that directly or indirectly influence dopamine signaling. Additional determinants of dopamine release include dopamine autoreceptors, muscarinic receptors, axon terminal density and intrinsic excitability (Troyano-Rodriguez et al., 2022; Arnsten et al., 1995; Alcantara et al., 2001; Shin et al., 2015; Razidlo et al., 2022; Voorn et al., 1988; Reynolds et al., 2025; Pokinko et al., 2017). Examining these mechanisms will further our understanding of age-dependent striatal function and corticostriatal organization.

Overall, we found normative striatal dopamine availability follows a chronologically similar lifespan trajectory across macaques and mice, with a prolonged period of increase between adolescence into adulthood. Species specific developmental changes in local GABAergic and cholinergic regulation contribute to, but cannot fully explain, this trajectory in each species. These results support the possibility that continual increases in dopamine release that coincide with, and perhaps facilitate, fronto-striatal circuit reorganization reflects a prolonged sensitive period. This work identifies mechanisms that contribute to the normative trajectory of striatal dopamine availability which can provide novel insights to understand age-dependent vulnerability to psychiatric disorders and may ultimately generate new targets for prevention and treatment.

## METHODS

*Animals:* All experiments were performed in accordance with the ILAR Guide for the Care and Use of Laboratory Animals and under an Animal Study Protocol approved by the ACUC at the National Institute of Mental Health and National Institute on Alcohol Abuse and Alcoholism. All procedures adhered to the applicable Federal and local laws, regulations, and standards, including the Animal Welfare Act and Regulations and Public Health Service policy (PHS2002). All animals were housed on a 12 h light/dark cycle (06:30 to 18:30 light) with food and water *ad libitum*. Macaque brain tissue was obtained from 22 adult rhesus macaques (*Macaca mulatta*, 10 male and 12 female; age 2–17 years, average age of 8.8 years), as part of the Comparative Brain Physiology Consortium at the National Institute of Health. Ten macaques were part of a larger project on adolescent development, while the remaining 12 were part of a larger project to test AAV delivery and expression in NHPs (Tangen et al., 2026); the data used in this manuscript were collected from non-injected hemispheres to prevent confounds. Mouse brain tissue was obtained from 59 C57BL/6J mice (27 male and 32 female; age 3-68 weeks, average age of 31 weeks). Sample sizes were limited and therefore cannot comprehensively resolve sex-dependent developmental effects, particularly in macaques. We also note that cross species comparisons are somewhat limited by regional striatal anatomy. Recordings sampled caudate and putamen in macaque and dorsomedial, but not dorsolateral, striatum in mouse.

To facilitate examination of cross-species patterns, data was binned into four lifespan periods defined by biological milestones shared across macaque and mouse. We identified four main periods: P1: prepuberty, P2: postpuberty, P3: postmaturity, and P4: postfertility. The average onset of puberty for macaques (2.5 years; Resko et al. 1982) and mice (4 weeks; Chester et al 2026) separated P1 and P2, skeletal maturity defined as bone plate closure in macaques (6.5 years; Trotter et al., 1975) and maximum bone density in C57BL/6mice (4 months; Jilka 2013) marked the beginning of P3, and the onset of declining fertility in mouse (50 weeks; Finch, 2014) the start of P4 in mice (**Fig 1c**). The onset of the post-fertile period coincides with the average lifespan in macaques, and therefore we could not, and did not include macaque data in a P4 period (Walker and Herndon, 2008).

*Brain extraction and slice preparation:* For macaque brain extraction, the animals were sedated with a mixture of ketamine and midazolam (ketamine 5–15 mg/kg, midazolam 0.05–0.3 mg/kg) and then maintained on isoflurane. A deep level of anesthesia was verified by an absence of response to toe-pinch and absence of response to corneal reflex. Prior to brain removal and blocking, macaques were transcardially perfused with ice-cold sucrose-substituted artificial cerebrospinal fluid (SSACSF) containing the following (in mM): 90 sucrose, 80 NaCl, 24 NaHCO3, 10 glucose, 3.5 KCl, 1.25 NaH2PO4, 4.5 MgCl2, and 0.5 CaCl2 saturated with 95% O2 and 5% CO2. Macaque brains were further sectioned to tissue blocks containing the caudate and putamen before thin (300 µm) sections were prepared. For mouse brain extraction, animals were anesthetized with isoflurane and brains quickly removed following decapitation and dropped into the same ice-cold SSACSF cutting solution before thin slices (240 µm) were prepared.

Macaque and mouse coronal brain slices were prepared using a Leica 1200 S Vibratome in the SSACSF solution saturated with 95% O2 and 5% CO2. Slices were incubated in a warm (∼32°C) SSACSF solution for 20–30 minutes and then returned to room temperature until recording. For recording, slices were transferred to a submerged recording chamber and perfused at 2 ml/min with extracellular artificial cerebrospinal fluid (ACSF) containing the following (in mM): 124 NaCl, 26.2 NaHCO3, 20 glucose, 2.5 KCl, 1 NaH2PO4, 2.5 CaCl2, 1.3 MgCl2, and 0.4 ascorbic acid saturated with 95% O2 and 5% CO2, which was heated at 32°C using an inline heater (Harvard Apparatus).

*Fast-scan cyclic voltammetry (FSCV):* FSCV was performed in freshly prepared coronal slices from the dorsal striatum in both species. Carbon fiber electrodes were prepared new each day with a cylindrical carbon fiber (7 µm in diameter; 230 µm of exposed fiber) inserted into a glass pipette, which was backfilled with 3 M KCl. Before use, the carbon fiber electrodes were conditioned with an 8-ms-long triangular voltage ramp (−0.4 to +1.2 and back to −0.4 V vs Ag/AgCl reference at 400 V/s) delivered every 15 ms for 20 minutes in the recording chamber filled with ACSF. The quality of each carbon-fiber electrode was assessed by measuring the peak current generated during the voltage ramp. Only data from electrodes with peak currents between 1.0 and 1.8 nA were included in the analysis.

After verifying electrode quality, the carbon fiber tip was inserted diagonally into the slice with a ∼22° angle to maximize contact between the carbon fiber surface and the tissue while minimizing the force pushing down into the tissue. During the recordings, carbon fiber electrodes were held at −0.4 V versus Ag/AgCl, and the same triangular voltage ramp was delivered every 100 ms. The stimulating electrode was positioned in close proximity to the stimulating electrode within the caudate or putamen (macaque) or the dorsal striatum (mouse). For electrical stimulation, a glass pipette filled with ACSF was placed near the tip of the carbon fiber and a brief rectangular pulse (0.2 ms for mouse slices; 2 ms for macaque slices; SIU91A by Cygnus Technologies) was applied every 2 minutes. When normalized to peak amplitude, voltammograms generated from evoked signals in mouse and macaque tissue closely overlapped and exhibited the characteristic oxidation and reduction peaks of dopamine at approximately +0.6 V and-0.2 V, respectively (**Extended Data Fig 1a**).

For input–output curves (**Extended Data Fig 1b**), the current amplitude for the stimulation electrode was adjusted to 10, 20, 40, 60, 100, 200, and 400 µA. Data were collected with a retrofit headstage (CB-7B/EC with 5 MΩ resistor) using a MultiClamp 700B amplifier (Molecular Devices) after low-pass filter at 3 kHz and digitized at 100 kHz using a digital-analog board (NI USB-6229 BNC, National Instruments). Data acquisition and analysis were performed using a custom-written software, VIGOR, in Igor Pro (WaveMetrics) using mafPC (courtesy of MA Xu-Friedman). The current peak amplitudes of the evoked dopamine transients were converted to dopamine concentration according to the calibration using 1–3 µM dopamine. Data from slices with highly variable amplitude of evoked dopamine signals, or unstable baseline current measurements, and from experiments with variable measurements of bath temperature were excluded from analyses.

*Measurement Calculations and Data Analysis:* In this study we report the following main measurements: maximum evoked dopamine and dopamine clearance as our primary measurements, alongside effects of pharmacological manipulation. We calculated maximum evoked dopamine for each brain slice by determining the maximum amplitude of the dopamine current trace (see Extended Data **Fig 1d** for examples) after the highest current stimulation (400mA, see **Extended Data Fig 1b**) and taking the average across five (5) baseline points in ACSF. For evoked dopamine, we used the same five (5) baseline transients to fit a single exponential decay function after the peak amplitude for each slice measurement and taking the average across those points in ACSF. For effects of pharmacology on dopamine release, we used a similar approach where after drugs were washed onto slices by being added to the perfusing ACSF (see Drugs and Chemicals for details), drug baseline points were averaged; for main figures, we calculated as a percentage of the ACSF baseline ((*drug baseline* / *ACSF baseline)*\*100), or by the change from baseline (100 - (*drug baseline* / *ACSF baseline)*\*100).

Data was organized and visualized in Python (*pandas, numpy, matplotlib, seaborn*) utilizing Jupyter notebooks. Statistical analysis was performed using R via an rpy2 bridge in the same Jupyter notebooks. The notebook and data are available via Mendeley Data. For each measurement, we examined macaque and mouse data separately, as our primary interest was age and age-period related changes over the lifespan.

For examining the relationship between age and dopamine measurements we found a generalized additive model (GAM) with a thin-plate regression spline (k = 4) (mgcv:gam()) with a restricted maximum likelihood best captured the non-linear nature of the data. We used a derivative analysis to determine peaks in each curve fit (gratia:derivatives()) to estimate where each species peaked in evoked dopamine. We used repeated measures ANOVA (stats:aov()) to examine the effect of lifespan period, sex, and recording region (for macaques) and period X sex interactions on dopamine release, dopamine clearance, and drug effects. Estimated marginal means tests were used following ANOVAs when appropriate with Tukey-adjusted p-values (emmeans:emmeans()). For cross-species comparisons, we limited the lifespan periods to P1, P2, and P3 as we did not have representative data in P4 for the macaques. We used rmAVONA to examine period, species, and period X species interactions with estimated marginal means for post-hoc analysis. For the results reported in this paper, we followed the field standard of treating each slice as an independent measurement. To corroborate these findings, we repeated these statistical analyses with rmANOVAs including an Error term for individual subjects to account for subject-to-subject variability and with linear mixed effects models with a random term for each subject and noted no major differences in the outcome of the analyses (**Extended Data Table**).

*Drugs and chemicals:* Dihydro-β-erythroidine hydrobromide (DHβE) and CGP55845 hydrochloride were purchased from Tocris, α-conotoxin-PIA from Alomone Labs, SR95531 (gabazine) from Hello Bio, and all other chemicals from Sigma-Aldrich. For drug application, we added each drug to the perfusing ACSF to achieve the following concentrations: Gabazine (5 µM) and CGP-55845 (2 µM); α-conotoxin-PIA (0.1 µM); α-conotoxin-PIA (0.1 µM) and DHβE (1µM). At this high concentration, DHβE has been shown to work as a general nAChR antagonist (Kirsch et al., 2016). For the Gabazine and CGP experiment, we washed on the drug for 30 minutes before collecting baseline values. For α-conotoxin-PIA, we washed on for 14 minutes and calculated the average of the last two points to examine the effect before proceeding with the combined α-conotoxin-PIA and DHβE wash; we waited 10 minutes after adding the DHβE before collecting the baseline points.

## ACKNOWLEDGEMENTS

The authors would like to thank all members of the Laboratory on the Neurobiology of Compulsive Behaviors and the Comparative Brain Physiology Consortium for critical feedback on this manuscript. We also thank Sebastian Peña-Vargas for his administrative support. The contributions of the NIH author(s) are considered Works of the United States Government. The findings and conclusions presented in this paper are those of the author(s) and do not necessarily reflect the views of the NIH or the U.S. Department of Health and Human Services.

## AUTHOR CONTRIBUTIONS

**Conceptualization:** S.R.W., M.A., V.A.A., and B.B.A.; **Experimental preparation:** macaque - M.A., P.H., A.C.T., K.A.P., R.A., A.C., J.H.S., H.C.G., and S.R.W.; mouse - J.H.S., R.R., H.C.G., and S.R.W.; **Experiment recordings:** macaque: S.R.W., J.H.S., H.C.G., E.S.S., R.R., A.L.C., L.A., and S.C.; mouse - S.R.W.; **Data Analysis:** S.R.W., B.B.A., and V.A.A.; **Writing - Original Draft:** S.R.W., and V.A.A.; **Review & Editing:** all authors; **Supervision:** V.A.A. and B.B.A.; **Resources:** V.A.A, B.B.A., and C.J.M.

## CODE AND DATA SHARING

Data will be available via Mendeley Data upon publication.

---

**Extended Data Fig. 1.**
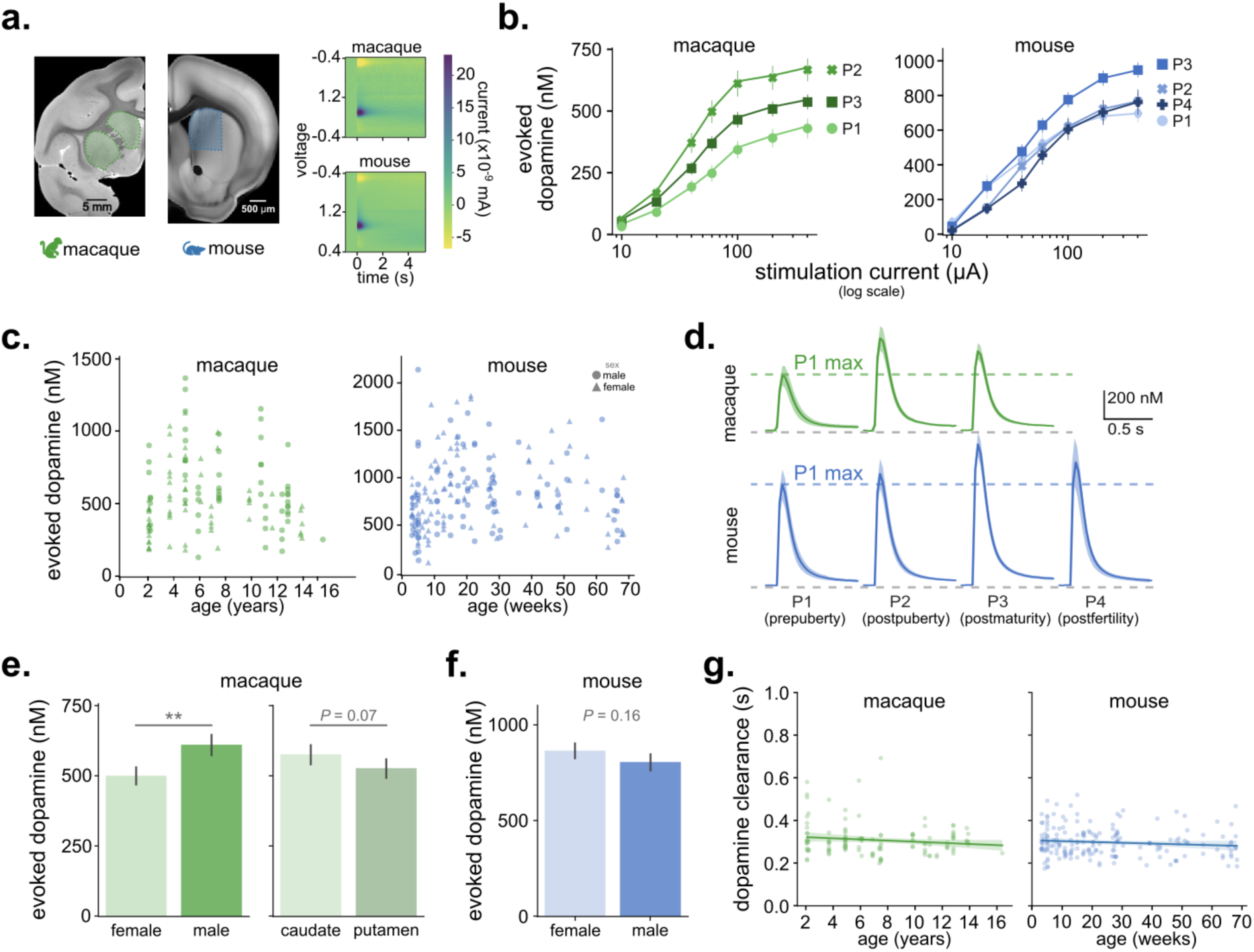
FSCV recording details and additional plots. a, Coronal hemisections of macaque and mouse brain showing recording locations in caudate and putamen (green) and dorsomedial striatum (blue). Representative FSCV color plots show current intensity as a function of time across the applied voltage ramp, with characteristic dopamine oxidation and reduction peaks at approximately +0.6 V and −0.2 V, respectively. **b,** Input-output curves between electrical stimulation intensity and evoked dopamine amplitude across lifespan periods. Symbols and lines show mean ± s.e.m. Evoked dopamine varied by lifespan period (macaque: F_2,854_ = 33.3, P < 0.001; mouse: F_3,1314_ = 14.5, P < 0.001) and stimulation intensity (macaque: F_1,854_ = 298.9, P < 0.001; mouse: F_1,1314_ = 479.5, P < 0.001), with no lifespan period × stimulation intensity interaction (macaque: F_2,854_ = 2.16, P = 0.115; mouse: F_3,1314_ = 1.84, P = 0.137). **c,** Maximum evoked dopamine per slice plotted against age. Females are shown as triangles and males as circles; shading progresses from light to dark across increasing lifespan periods. **d,** Average evoked dopamine transients by lifespan period in macaque and mouse striatum. Lines and shading show mean ± 95% confidence interval. **e,** Mean evoked dopamine in macaques by sex (left) and striatal region (right). Release was lower in females than males (F_1,119_ = 9.35, P = 0.003) and showed a trend toward greater release in caudate than putamen (F_1,119_ = 3.17, P = 0.07). **f,** Mean evoked dopamine in mice by sex. Release did not differ between males and females (F_1,181_ = 1.99, P = 0.16). **g,** Dopamine transient decay time constant (τ) plotted against age. Solid lines show linear fits with 95% confidence intervals. τ was not significantly associated with age in macaques (β = −0.002, s.e. = 0.001, t_125_ = −1.61, P = 0.11; linear model) or mice (β = −0.0003, s.e. = 0.0002, t_125_ = −1.51, P = 0.134; linear model). *Repeated-measures ANOVA was used unless otherwise indicated. Error bars represent mean ± s.e.m. Post hoc comparisons were performed using estimated marginal means with Tukey adjustment for multiple comparisons. *P < 0.05, **P < 0.01, ***P < 0.001.*

**Extended Data Fig. 2.**
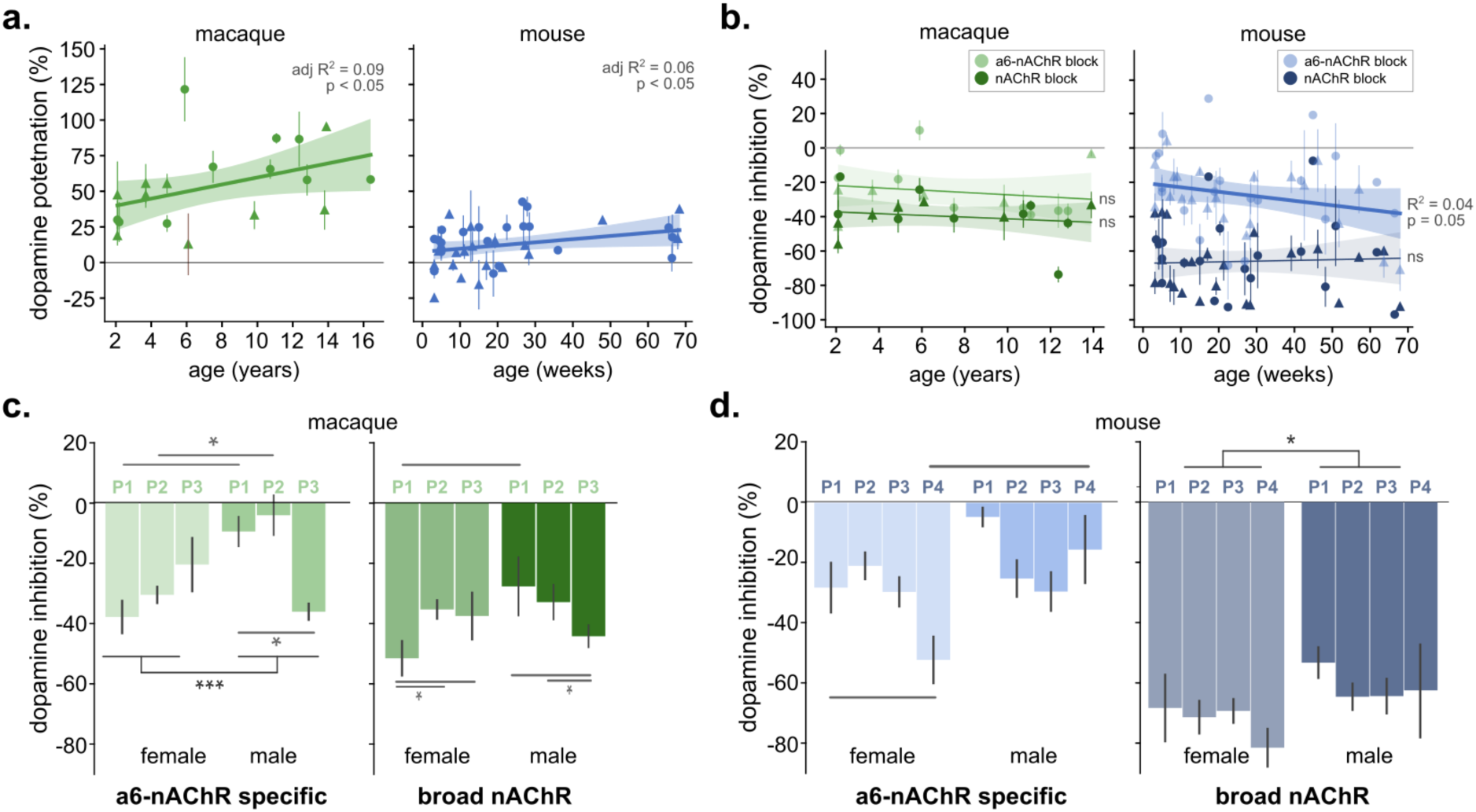
Additional analyses of pharmacological impact on dopamine release regulation. a, Percent change in evoked dopamine following GABA receptor antagonism plotted against age. Solid lines show linear fits with shaded 95% confidence intervals; points show mean effects per subject (macaque: N = 19, n = 57; mouse: N = 40, n = 78). GABA receptor antagonist-induced potentiation increased with age in macaques (β = 2.06, s.e. = 0.97, t_53_ = 2.12, P = 0.039) and mice (β = 0.189, s.e. = 0.094, t_75_ = 2.02, P = 0.047). In macaques, effects did not differ between caudate and putamen (β = −5.99, s.e. = 7.98, t_53_ = −0.75, P = 0.456) or by sex (β = 11.1, s.e. = 8.30, t_53_ = 1.34, P = 0.18). No sex effect was detected in mice (β = 6.01, s.e. = 3.82, t_75_ = 1.58, P = 0.120). **b,** Percent change in evoked dopamine following nAChR antagonism plotted against age. Solid lines show linear fits with shaded 95% confidence intervals; points show mean effects per subject (macaque: N = 16, n = 56; mouse: N = 43, n = 98). Lighter shades denote α6- nAChR blockade and darker shades denote broader nAChR blockade. In macaques, effects did not differ between caudate and putamen (β = −1.58, s.e. = 4.93, t_49_ = −0.32, P = 0.750). The effect of α6-nAChR blockade was not associated with age (β = −1.03, s.e. = 0.66, t_49_ = −1.56, P = 0.125), nor was the effect of broader nAChR blockade (β = −0.49, s.e. = 0.60, t_48_ = −0.83, P = 0.412). In mice, α6-nAChR antagonist effects increased in magnitude with age (β = −0.311, s.e. = 0.13, t_86_ = −2.42, P = 0.018), whereas broader nAChR antagonist effects were not associated with age (β = −0.093, s.e. = 0.12, t_77_ = −0.77, P = 0.443). **c,** Effects of α6-nAChR blockade (left) and broader nAChR blockade (right) by sex and lifespan period in macaques. α6-nAChR antagonist effects differed by sex (F_1,46_ = 8.46, P = 0.006) and showed a period × sex interaction (F_2,46_ = 12.4, P < 0.001; see Extended Data Table for post hoc comparisons). Broader nAChR antagonist effects also showed a period × sex interaction (F_2,45_ = 4.16, P = 0.022). **d,** Effects of α6-nAChR blockade (left) and broader nAChR blockade (right) by sex and lifespan period in mice. α6-nAChR antagonist effects did not differ by sex (F_1,81_ = 0.99, P = 0.32), with a trend toward a period × sex interaction (F_3,81_ = 2.63, P = 0.056). Broader nAChR antagonist effects showed neither a main effect of sex (F_1,72_ = 2.93, P = 0.09) nor a period × sex interaction (F_3,72_ = 0.34, P = 0.79). *Repeated-measures ANOVA was used unless otherwise indicated. Error bars represent mean ± s.e.m. Post hoc comparisons were performed using estimated marginal means with Tukey adjustment for multiple comparisons. *P < 0.05, **P < 0.01, ***P < 0.001.*

